# Fission yeast Rad8/HLTF facilitates Rad52-dependent chromosomal rearrangements through PCNA lysine 107 ubiquitination

**DOI:** 10.1101/2021.01.05.425428

**Authors:** Jie Su, Naoko Toyofuku, Takuro Nakagawa

## Abstract

Rad52 recombinase can cause gross chromosomal rearrangements (GCRs). However, the mechanism of Rad52-dependent GCRs remains unclear. Here, we show that fission yeast Rad8/HLTF facilitates Rad52-dependent GCRs through the ubiquitination of lysine 107 (K107) of PCNA, a DNA sliding clamp. Loss of Rad8 reduced isochromosomes resulting from centromere inverted repeat recombination. Rad8 HIRAN and RING finger mutations reduced GCRs, suggesting that Rad8 facilitates GCRs through 3’ DNA-end binding and ubiquitin ligase activity. Mms2 and Ubc4 but not Ubc13 ubiquitin-conjugating proteins were required for GCRs. Consistent with this, PCNA K107R but not K164R mutation greatly reduced GCRs. Rad8-dependent PCNA K107 ubiquitination facilitates Rad52-dependent GCRs, as PCNA K107R, *rad8*, and *rad52* mutations epistatically reduced GCRs. Remarkably, K107 is located at the interface between PCNA subunits, and an interface mutation D150E bypassed the requirement of PCNA K107 ubiquitination for GCRs. This study uncovers the role of Rad8-dependent PCNA K107 ubiquitination in Rad52-dependent GCRs.

## Introduction

Faithful repair of DNA damage, such as DNA double-strand breaks (DSBs), is critical to maintaining genome integrity *(Jeggo et al., 2016; Putnam & Kolodner, 2017)*. Homologous recombination is considered an error-free DSB repair mechanism, as it uses intact DNA as the template. However, when accompanied by crossover or break-induced replication, nonallelic homologous recombination between repetitive sequences in a genome results in gross chromosomal rearrangements (GCRs), including translocation, deletion, and inversion *(Carvalho & Lupski, 2016; Symington et al., 2014)*. Isochromosomes, whose arms mirror each other, are the GCR product formed by recombination between inverted repeats around centromeres *(Nakagawa & Okita, 2019)*. GCRs can cause cell death and genetic diseases, including cancer *(Weischenfeldt et al., 2013)*. Therefore, it is important to elucidate how GCRs occur for preventing those diseases.

Rad51 and Rad52 are evolutionally conserved recombination enzymes that have pivotal roles in distinct recombination pathways *(Bennardo et al., 2008; Chang et al., 2017)*. Rad51 preferentially promotes conservative recombination, gene conversion *(Onaka et al., 2016; Rattray & Symington, 1994; Stark et al., 2004)*. Rad51 binds single-stranded DNA (ssDNA) and catalyzes DNA strand exchange with homologous double-stranded DNA (dsDNA) *(Kowalczykowski, 2015)*. Mammalian BRCA1 and BRCA2 facilitate Rad51-dependent recombination. Mutation in BRCA genes increases GCRs and predisposes the carriers to hereditary breast and ovarian cancer *(Nielsen et al., 2016)*, demonstrating that Rad51-dependent recombination safeguards genome integrity and prevents tumorigenesis. Although yeast Rad52 mediates Rad51 loading onto RPA-coated ssDNA, human Rad52 has no such activity, and BRCA2 mediates Rad51 loading *(Kowalczykowski, 2015)*. In contrast to the mediator function, both yeast and human Rad52 proteins catalyze single-strand annealing (SSA) by which complementary ssDNA strands are annealed to form dsDNA *(Kagawa et al., 2002; Mortensen et al., 1996; Reddy et al., 1997)* and RNA or DNA strand exchange with homologous dsDNA *(Kagawa et al., 2001; Mazina et al., 2017)*. For the sake of simplicity, Rad52-dependent SSA and strand exchange that occur independently of Rad51 are designated Rad52-dependent recombination in this paper. Rad52 knockout mice show only a mild defect in DNA recombinational repair and are not predisposed to cancer *(Rijkers et al., 1998)*. However, Rad52 deficiency is synthetic lethal with BRCA mutations *(Hanamshet et al., 2016; Jalan et al., 2019)*. Rad52 N-terminal domain that retains SSA and strand exchange activities is sufficient to restore cell growth of the double mutants *(Hanamshet & Mazin, 2020)*, showing the importance of Rad52-dependent recombination in cells defective in Rad51-dependent recombination. Consistent with this idea, yeast *rad52*Δ cells exhibit more severe DNA repair and recombination defects than *rad51*Δ cells *(Symington et al., 2014)*. Previously, we have shown in fission yeast that loss of Rad51 reduces gene conversion between centromere inverted repeats and increases isochromosome formation in a manner dependent on Mus81 crossover-specific endonuclease *(Nakamura et al., 2008; Onaka et al., 2016)*. The *rad52-R45K* mutation in the N-terminal domain impairs *in vitro* SSA activity and reduces isochromosome formation in *rad51*Δ cells *(Onaka et al., 2020)*, showing that Rad52-dependent recombination promotes isochromosome formation. At fission yeast centromeres, Rad51-dependent recombination predominates, and Rad52-dependent recombination is suppressed *(Zafar et al., 2017)*. Intriguingly, a mutation in DNA polymerase α (Pol α) required for lagging-strand synthesis increases Rad52-dependent recombination without changing the total rate of recombination at centromeres *(Onaka et al., 2020)*, suggesting that the formation of ssDNA gaps during DNA replication might be a rate-limiting step of Rad52-dependent recombination. Rad52-dependent recombination leads not only GCRs but also gene conversion in the absence of Rad51. Msh2-Msh3 MutS-homologs *(Surtees & Alani, 2006)* and Mus81 crossover-specific endonuclease *(Doe et al., 2004; Osman et al., 2003)* are required for the GCR but not gene conversion branch of Rad52-dependent recombination that occurs in the absence of Rad51 *(Onaka et al., 2020; Onaka et al., 2016)*, suggesting that Msh2-Msh3 and Mus81 resolve joint molecules formed by Rad52 specifically into GCR products. However, how Rad52-dependent recombination differentiates into GCRs is still unclear.

Proliferating cell nuclear antigen (PCNA) protein interacts with each other and forms a homotrimeric DNA clamp that serves as a landing pad for the factors related to replication and repair *(Krishna et al., 1994; Moldovan et al., 2007)*. Replication factor C (RFC) complexes load the PCNA clamp onto DNA at primer ends *(Majka & Burgers, 2004)*. After ligation of Okazaki fragments, RFC-like complexes containing Elg1 unload PCNA from DNA *(Kubota et al., 2015; Kubota et al., 2013)*. Post-translational modifications of PCNA are critical in regulating its function *(Leung et al., 2018)*. Among them, the ubiquitination of PCNA has been most well studied *(Hoege et al., 2002)*. Rad18 ubiquitin ligase (E3) and Rad6 ubiquitin-conjugating enzyme (E2) catalyze PCNA K164 mono-ubiquitination to recruit translesion synthesis polymerases, including DNA Pol η. Depending on the mono-ubiquitination, budding yeast Rad5 (E3) and Mms2-Ubc13 (E2) catalyze K164 poly-ubiquitination to promote template switching, a recombination-mediated damage bypass pathway *(Giannattasio et al., 2014)*. Fission yeast Rad8 and human HLTF are homologs of budding yeast Rad5. They are unique among ubiquitin ligases as they contain HIRAN and SNF/SWI helicase domains besides the E2 binding domain RING finger *(Frampton et al., 2006; Hoege et al., 2002; Motegi et al., 2008)*. HIRAN is a modified oligonucleotide/oligosaccharide (OB) fold domain that binds 3’ DNA-ends *(Achar et al., 2015; Hishiki et al., 2015; Kile et al., 2015)*. Interestingly, in the *cdc9* mutant strain of DNA ligase I essential for Okazaki fragment ligation, budding yeast Rad5 (E3) ubiquitinates PCNA at K107 rather than K164 *(Das-Bradoo et al., 2010; Nguyen et al., 2013)*. K107 ubiquitination requires Mms2-Ubc4 (E2) but not Mms2-Ubc13 (E2) and occurs independently of Rad18 (E3) or Rad6 (E2). K107 ubiquitination is required for the full activation of a checkpoint kinase in *cdc9* mutants. However, it is unknown how K107 ubiquitination affects PCNA structure and function.

Here, we found that fission yeast Rad8 (E3) and Mms2-Ubc4 (E2) ubiquitinate PCNA at K107 to facilitate Rad52-dependent GCRs. Loss of Rad8 reduced isochromosome formation but not chromosomal truncation in *rad51*Δ cells. Mutation in Rad8 HIRAN or RING finger but not helicase domain reduced GCRs. *mms2* and *ubc4* but not *ubc13*; PCNA K107R but not K164R reduced GCRs in *rad51*Δ cells, indicating that the ubiquitination of K107 but not a canonical site K164 is required for GCRs. The epistatic analysis showed that Rad8-dependent PCNA K107 ubiquitination plays a crucial role in the GCR branch of Rad52-dependent recombination. K107 is located at the interface between PCNA subunits, suggesting that K107 ubiquitination weakens the PCNA-PCNA interaction to cause GCRs. Remarkably, an interface mutation D150E *(Goellner et al., 2014; Johnson et al., 2016)* indeed bypassed the requirement of K107 ubiquitination for GCRs. These data suggest that PCNA K107 ubiquitination weakens the subunit interaction of the PCNA clamp to facilitate Rad52-dependent GCRs.

## Results

### Fission yeast Rad8 facilitates isochromosome formation but not chromosomal truncation

Loss of Rad8 increases DNA damage sensitivity of *rad51*Δ cells *(Ding & Forsburg, 2014; Frampton et al., 2006)*, raising the possibility that Rad8 is involved in GCRs that occur independently of Rad51. To test this, we disrupted the *rad8* gene in the fission yeast cell harbouring the extra-chromosome ChL^C^ derived from chromosome 3 (chr3) (Figure 1A). ChL^C^ retains an entire region of centromere 3 (cen3) that consists of a unique sequence, cnt3, surrounded by inverted repeats: imr3, dg, dh, and irc3, and contains three genetic markers: *LEU2, ura4*^+^, and *ade6*^+^ *(Nakamura et al., 2008; Onaka et al., 2016)*. As ChL^C^ is dispensable for cell growth, one can detect otherwise lethal GCRs using ChL^C^. To detect Leu^+^ Ura^−^ Ade^−^ GCR clones that have lost both *ura4*^+^ and *ade6*^+^, cells were grown in Edinburgh minimum media supplemented with uracil and adenine (EMM+UA) and plated on FOA+UA media containing 5-fluoroorotic acid (5-FOA) which is toxic to *ura4*^+^ cells. Leu^+^ Ura^−^ colonies formed on FOA+UA plates were transferred to EMM+U to inspect adenine auxotrophy. Fluctuation analysis showed that *rad8*Δ did not significantly change spontaneous GCR rates in wild-type cells (Figure 1B). However, *rad8*Δ partially but significantly reduced GCRs in *rad51*Δ cells (Figure 1B), indicating that Rad8 facilitates GCRs that occur independently of Rad51.

**Figure 1.**
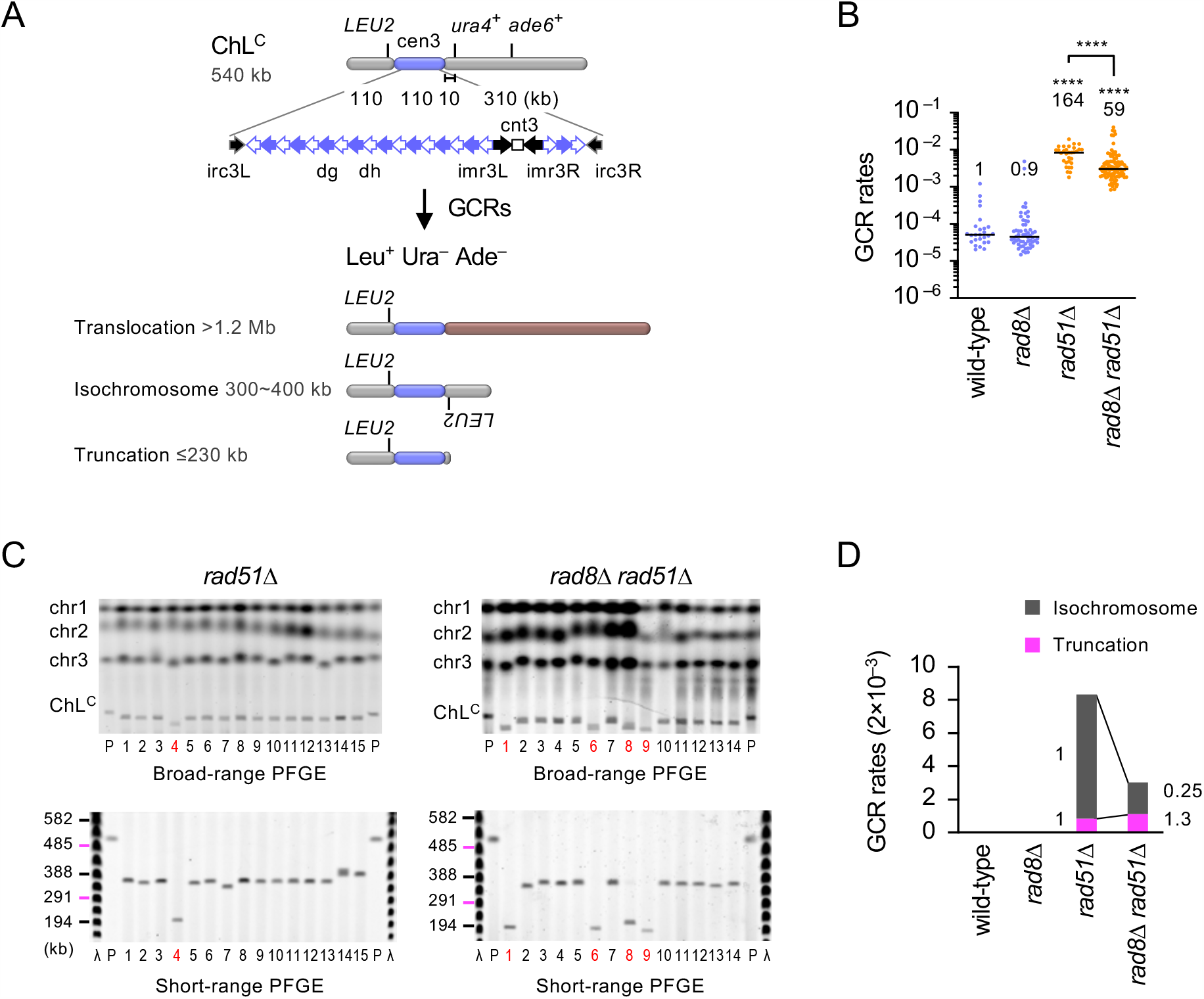
Fission yeast Rad8 facilitates isochromosome formation. (**A**) The ChL^C^ chromosome contains an entire region of centromere 3 (cen3) and three genetic markers: *LEU2, ura4*^+^, and *ade6*^+^. Gross chromosomal rearrangements (GCRs) associated with loss of both *ura4*^+^ and *ade6*^+^ result in the formation of Leu^+^ Ura^−^ Ade^−^ cells. The structure and the length of three GCR types: translocation, isochromosome, and truncation are shown. (**B**) GCR rates of the wild-type, *rad8*Δ, *rad51*Δ, and *rad8*Δ *rad51*Δ strains (TNF5369, 5549, 5411, and 5644, respectively). Each dot represents an independent experimental value obtained from an independent colony. Black bars indicate medians. Rates relative to that of wild type are shown on the top of each cluster of dots. Statistical analyses between the wild-type and mutant strains, and that between *rad51*Δ and *rad8*Δ *rad51*Δ strains were performed by the two-tailed Mann-Whitney test. **** *P* < 0.0001. (**C**) Chromosomal DNAs prepared from the parental (P) and independent GCR clones of *rad51*Δ and *rad8*Δ *rad51*Δ strains were separated by broad- and short-range PFGE (top and bottom rows, respectively). Positions of chr1, chr2, chr3, and ChL^C^ (5.7, 4.6, ∼3.5, and 0.5 Mb, respectively) are indicated on the left of broad-range PFGE panels. Sizes of Lambda (λ) ladders (ProMega-Markers, Madison, Wisconsin, G3011) are displayed on the left of short-range PFGE panels. Sample number of truncations are highlighted in red. GCR products for wild-type and *rad8*Δ and additional samples for *rad51*Δ and *rad8*Δ *rad51*Δ strains are shown in Figure 1—figure supplement 1. (**D**) Rates of each GCR type. Rates relative to those of *rad51*Δ are indicated. Source data of the graphs and uncropped images of the gels are available in Figure 1—Source Data 1 and 2, respectively.

Three types of GCRs have been detected using ChL^C^: translocation, isochromosome, and truncation, which can be distinguished by the length *(Nakamura et al., 2008; Onaka et al., 2020; Onaka et al., 2016)* (Figure 1A). Among them, isochromosomes are formed by recombination between inverted repeats at centromeres. To determine GCR types Rad8 causes, chromosomal DNAs were prepared from the parental and independent GCR clones, resolved by pulsed-field gel electrophoresis (PFGE) under two different conditions (broad- and short-range PFGE), and stained with ethidium bromide (EtBr) (Figure 1C; results for wild-type and *rad8*Δ and additional results for *rad51*Δ and *rad8 rad51*Δ are in Figure 1—figure supplement 1). As previously observed *(Onaka et al., 2020)*, isochromosomes and a small number of translocations were detected in wild-type cells (Table 1). *rad8*Δ did not significantly change the GCR types in wild-type cells (*P* = 0.6, the two-tailed Fisher’s exact test). As previously observed *(Onaka et al., 2020), rad51*Δ cells produced isochromosomes and a small number of truncations (Figure 1C, sample #4). Given elevated GCR rates (Figure 1B), these results show that *rad51*Δ increases isochromosome formation and chromosomal truncation. In *rad51*Δ cells, *rad8*Δ increased the proportion of truncations from 10 to 37% (Table1; *P* = 0.030, the two-tailed Fisher’s exact test). To obtain the rate of each GCR type (Figure 1D), we multiplied the total GCR rate (Figure 1B) by the proportion of each GCR type (Table 1). Figure 1D shows that Rad8 is required for 75% of isochromosomes produced in *rad51*Δ cells. However, Rad8 is dispensable for chromosomal truncation.

**Table 1.** GCR types.

|  | translocation | isochromosome | truncation* | total |
| --- | --- | --- | --- | --- |
| wild type | 1 (3%) | 31 (97%) | 0 [0] | 32 |
| *rad8*Δ | 2 (7%) | 28 (93%) | 0 [0] | 30 |
| *rad51*Δ | 0 | 27 (90%) | 3 [2] (10%) | 30 |
| *rad8*Δ *rad51*Δ | 0 | 19 (63%) | 11 [9] (37%) | 30 |

Percentages of each GCR type are shown in ( ).
\*The numbers of the truncation products whose breakpoints are present in centromere repeats are shown in [ ].

It has been shown that truncation ends are healed by *de novo* telomere addition (Dave et al., 2020; Matsumoto et al., 1987; Pennaneach et al., 2006). To determine the chromosomal sites to which telomere sequences have been added, we amplified the breakpoints using a pair of ChL^C^ and telomere primers: M13R-C19 or M13R-T1 *(Irie et al., 2019)* and chromosomal truncations recovered from agarose gels. DNA sequencing of the PCR products revealed that telomere sequences (G_2-5_TTAC (A) (C)) *(Hiraoka et al., 1998)* were added either within or outside centromere repeats (Table 1 and 2). Only 0∼3-bp overlaps were detected between ChL^C^ and telomere sequences around the breakpoints, suggesting that no extensive annealing with telomere RNA is required to initiate telomere addition. No apparent differences were detected between *rad51*Δ and *rad8*Δ *rad51*Δ. Together, these results show that Rad8 is required for homology-mediated GCRs resulting in isochromosome formation but dispensable for *de novo* telomere addition resulting in chromosomal truncation.

**Table 2.**
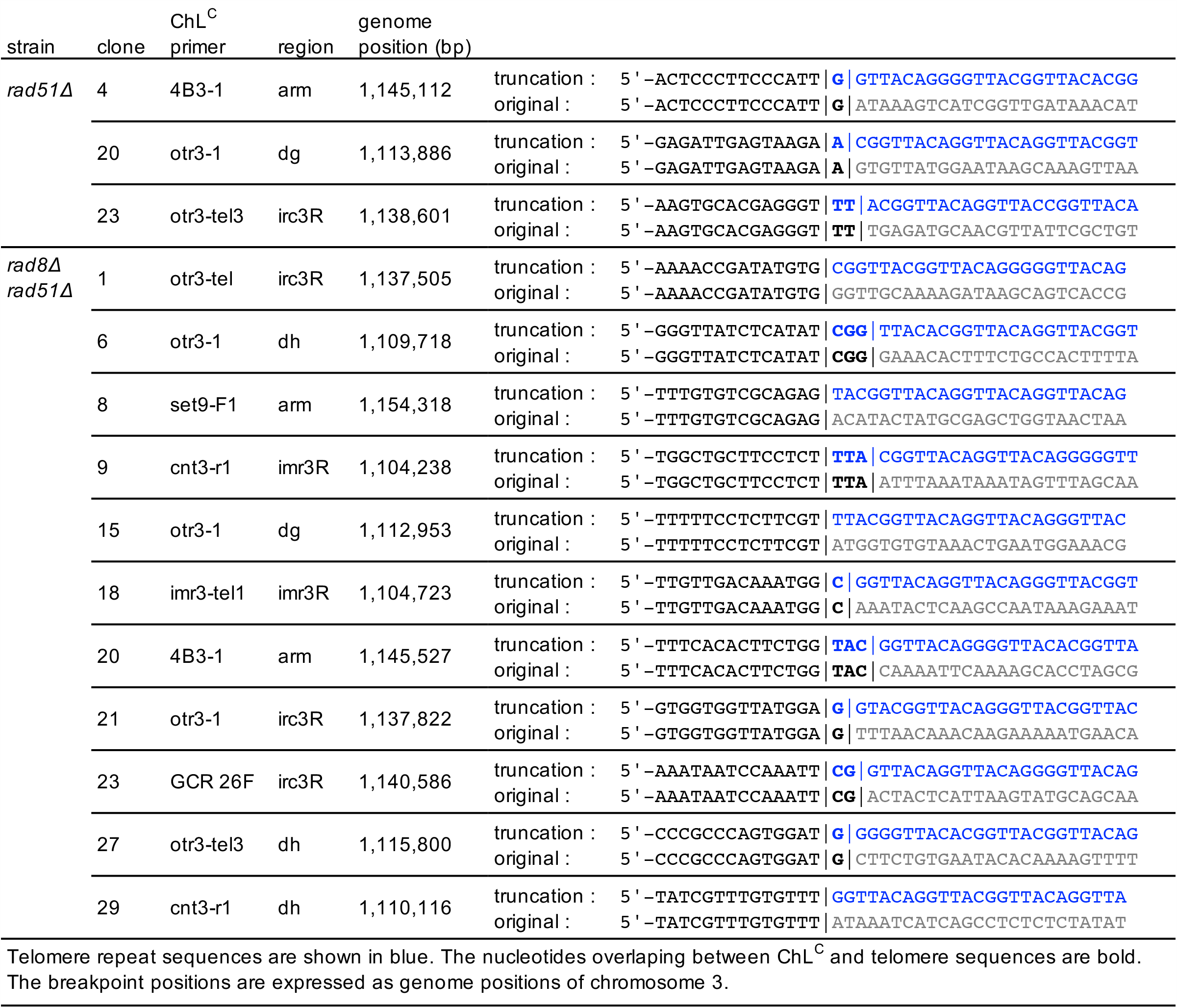
Breakpoints of chromosomal truncations

### Rad8 HIRAN and RING finger domains are required for GCRs

Rad8 and its homologs contain HIRAN, helicase, and RING finger domains (Figure 2A, top). To gain insight into how Rad8 facilitates GCRs, we introduced alanine (A) substitution into each domain (Figure 2A, bottom). The *rad8-HIRAN* mutation alters the residues forming the 3’ DNA-end binding pocket *(Achar et al., 2015; Hishiki et al., 2015; Kile et al., 2015)*. The *rad8-Helicase* mutation alters the conserved residues in the ATP-binding Walker A motif *(Blastyak et al., 2010)*. The *rad8-RING* mutation changes a residue involved in the interaction with ubiquitin-conjugating enzymes *(Ulrich, 2003; Ulrich & Jentsch, 2000). rad8-Helicase* did not significantly change GCR rates in both wild-type and *rad51*Δ cells, indicating that Rad8 helicase activity is not essential for GCRs. On the other hand, *rad8-HIRAN* and *rad8-RING* reduced GCR rates (Figure 2B), suggesting that Rad8 facilitates GCRs through 3’ DNA-end binding and ubiquitin ligase activity. The mutant proteins may have dominant-negative effects on GCRs, as *rad8-HIRAN* and *rad8-RING* exhibited slightly more pronounced effects on GCR rates than *rad8*Δ (Figures 1B and 2B).

**Figure 2.**
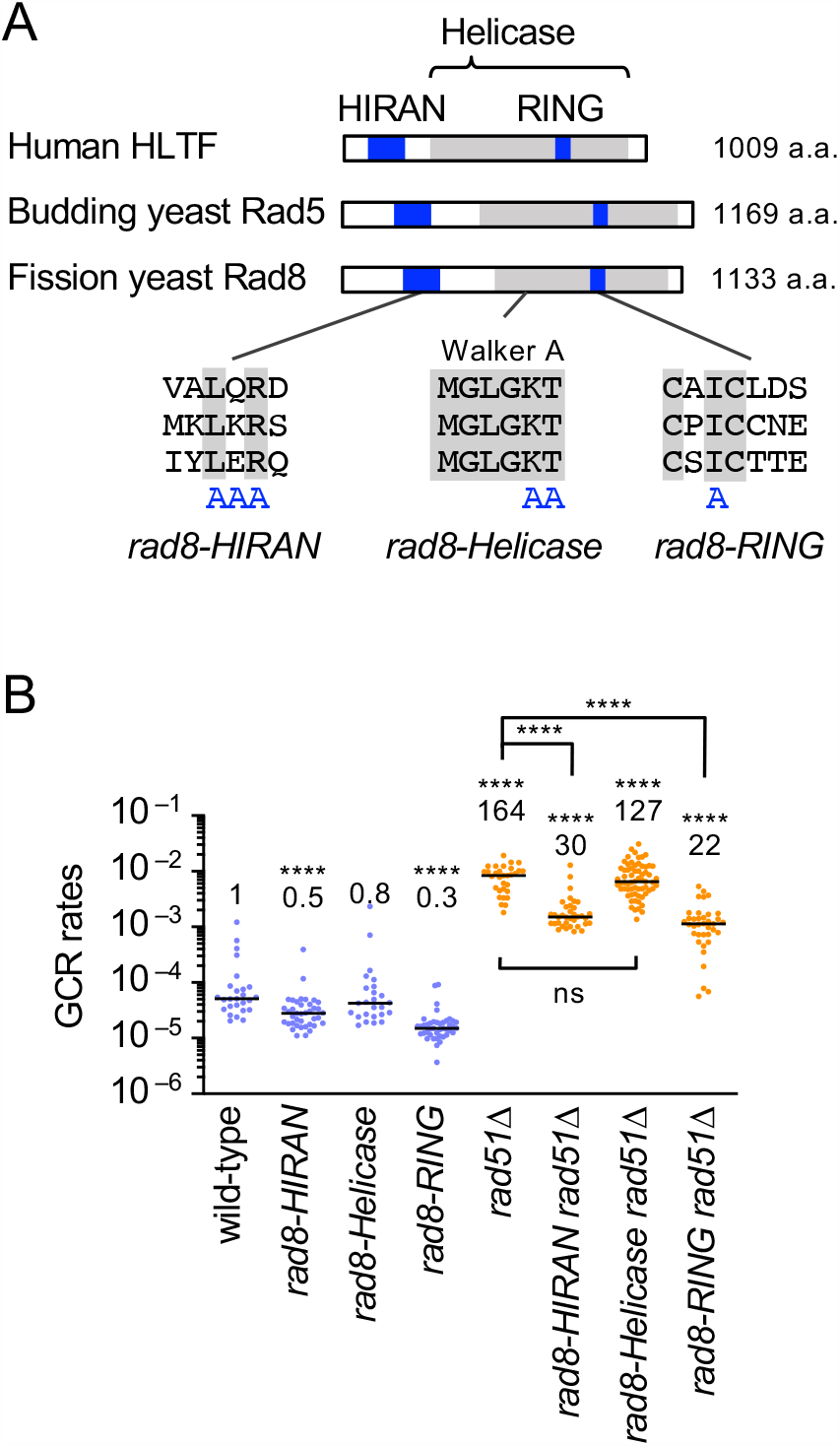
Rad8 HIRAN and RING domains are required for GCRs. (**A**) A schematic diagram showing the HIRAN, Helicase, and RING domains of fission yeast Rad8, budding yeast Rad5, and human HLTF. Amino acid residues substituted for alanine in the *rad8-HIRAN, rad8-Helicase*, and *rad8-RING* mutations are indicated. (**B**) GCR rates of the wild-type, *rad8-HIRAN, rad8-Helicase, rad8-RING, rad51*Δ, *rad8-HIRAN rad51*Δ, *rad8-Helicase rad51*Δ, and *rad8-RING rad51*Δ strains (TNF5369, 6205, 6203, 6207, 5411, 6217, 6231, and 6219, respectively). The two-tailed Mann-Whitney test. Non-significant (ns) *P* > 0.05; **** *P* < 0.0001. Source data of the graph are available in Figure 2—Source Data 1.

### Rad8 facilitates GCRs through PCNA K107 ubiquitination with the aid of Mms2-Ubc4

Rad8 ubiquitinates PCNA at K164 with the aid of Mms2-Ubc13 ubiquitin-conjugating complex, to promote template switching *(Frampton et al., 2006)* (Figure 3A). Rad8 also ubiquitinates PCNA at K107 with the aid of Mms2-Ubc4 complex, by inference from excellent studies of budding yeast Rad5 *(Das-Bradoo et al., 2010; Nguyen et al., 2013)*. To determine whether Rad8 facilitates GCRs with these ubiquitin-conjugating enzymes, we constructed *mms2, ubc13*, and *ubc4* mutant strains. As *ubc4* is an essential gene, we created the *ubc4-P61S* point mutation that impairs protein ubiquitination *(Seino et al., 2003). mms2*Δ reduced GCRs in *rad51*Δ cells (Figure 3B). To our surprise, *ubc13*Δ did not significantly change GCR rates and GCR types (Figure 3—figure supplement 1), but *ubc4-P61S* reduced GCRs in *rad51*Δ cells. It should be noted that the *rad8-RING* mutation did not further reduce GCRs in *ubc4-P61S rad51*Δ cells. These results show that Rad8 facilitates GCRs with the aid of Mms2-Ubc4 rather than Mms2-Ubc13.

**Figure 3.**
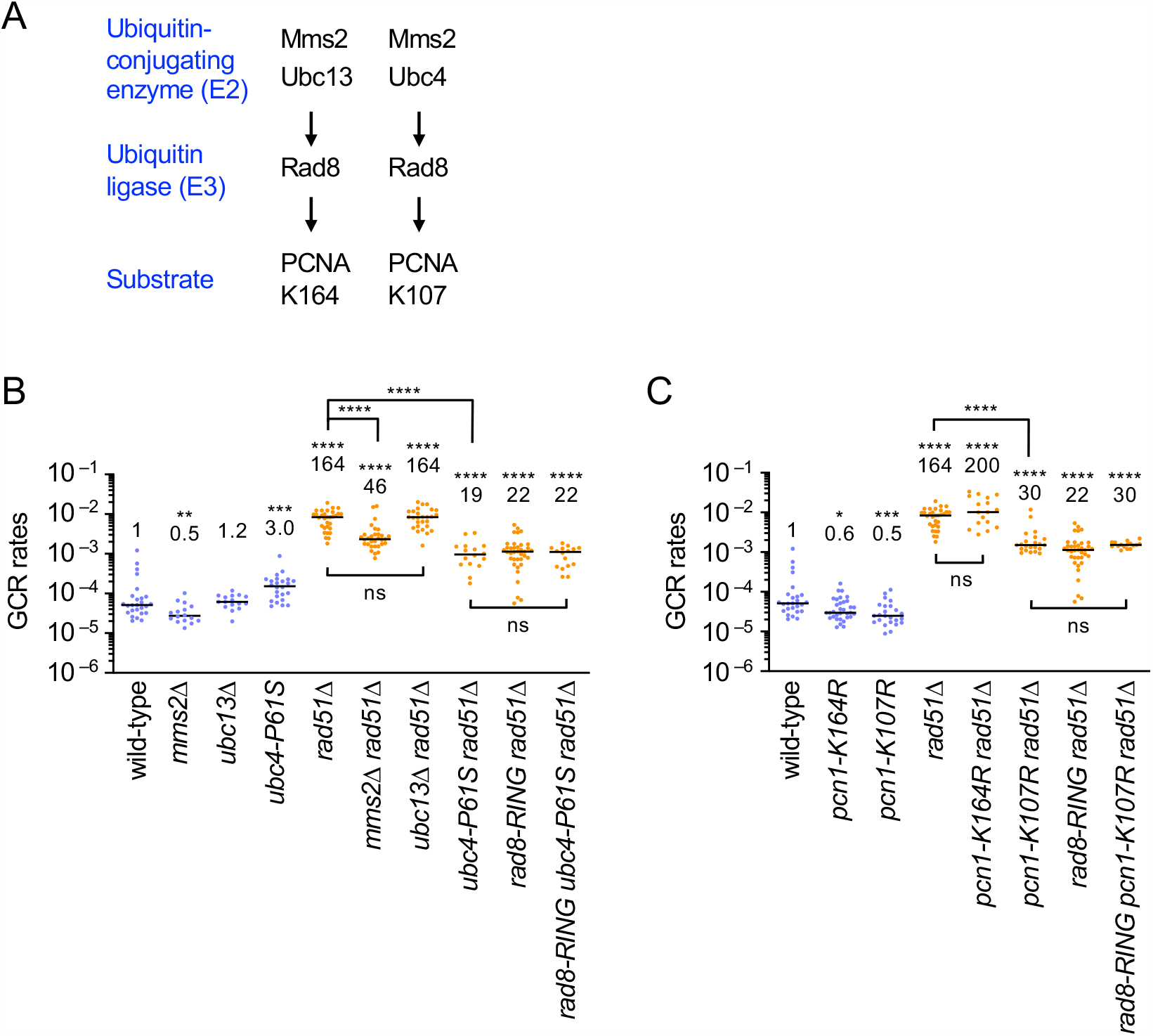
With the aid of Mms2-Ubc4, Rad8 ubiquitinates PCNA at K107 to cause GCRs. (**A**) Two distinct PCNA ubiquitination pathways. Human HLTF, budding yeast Rad5, and fission yeast Rad8 ubiquitinate PCNA K164 with the aid of Mms2-Ubc13 complex. Rad5 ubiquitinates PCNA K107 with the aid of the Mms2-Ubc4 complex. (**B**) GCR rates of the wild-type, *mms2*Δ, *ubc13*Δ, ubc4-P61S, *rad51*Δ, *mms2*Δ *rad51*Δ, *ubc13*Δ *rad51*Δ, *ubc4*-*P61S rad51*Δ, *rad8-RING rad51*Δ, and *rad8-RING ubc4-P61S rad51*Δ strains (TNF5369, 6751, 5915, 7484, 5411, 6771, 6115, 7503, 6219, and 7501, respectively). (**C**) GCR rates of the wild-type, *pcn1-K164R, pcn1-K107R, rad51*Δ, *pcn1-K164R rad51*Δ, *pcn1-K107R rad51*Δ, *rad8-RING rad51*Δ, and *rad8-RING pcn1-K107R rad51*Δ strains (TNF5369, 6078, 6738, 5411, 6104, 6761, 6219, and 6999, respectively). The two-tailed Mann-Whitney test. Non-significant (ns) *P* > 0.05; * *P* < 0.05; ** *P* < 0.01; *** *P* < 0.001; **** *P* < 0.0001. GCR products formed in *ubc13*Δ *rad51*Δ and *pcn1-K164R rad51*Δ cells are shown in Figure 3—figure supplement 1. GCR rates of *pcn1-K107A* strains are shown in Figure 3—figure supplement 2. DNA damage sensitivities of *pcn1-K164R* and *pcn1-K107R* strains are shown in Figure 3—figure supplement 3. Rad52 focus formation in *pcn1-K107R* strains is shown in Figure 3—figure supplement 4. Source data of the graphs are available in Figure 3—Source Data 1

The data presented above suggest that the ubiquitination of PCNA K107 rather than the well-known K164 is involved in GCRs (Figure 3A). To test this, we replaced the lysine (K) residue with arginine (R), to which no ubiquitins are conjugated, and determined GCR rates of the *pcn1* mutant strains (Figure 3C, the *pcn1* gene encodes PCNA). In wild-type cells, both *pcn1-K107R* and *pcn1-K164R* slightly reduced GCR rates (see Discussion). However, only *pcn1-K107R* reduced GCRs in *rad51*Δ cells (Figure 3C and Figure 3—figure supplement 1). Like *pcn1-K107R, pcn1-K107A* reduced GCRs (Figure 3—figure supplement 2), demonstrating the importance of the ubiquitin acceptor lysine in GCRs. Notably, the *rad8-RING* mutation did not further reduce GCRs in *pcn1-K107R rad51*Δ cells. Together, these results show that, with the aid of the Mms2-Ubc4 ubiquitin-conjugating complex, Rad8 ubiquitin ligase ubiquitinates PCNA K107 to facilitate GCRs.

### Rad8-dependent PCNA K107 ubiquitination is involved in the Rad52-dependent GCR pathway

We have shown that Rad52 causes isochromosome formation in *rad51*Δ cells, using the *rad52-R45K* mutation that impairs SSA activity of Rad52 protein *(Onaka et al., 2020)*. To determine whether Rad8-dependent PCNA K107 ubiquitination plays a role in the Rad52-dependent GCR pathway, we introduced the *rad8*Δ or *pcn1-K107R* mutation into *rad52-R45K rad51*Δ cells (Figure 4A). As previously observed, *rad52-R45K rad51*Δ cells showed reduced GCR rates than *rad51*Δ cells. Neither *rad8*Δ nor *pcn1-K107R* significantly reduced GCRs in *rad52-R45K rad51*Δ cells. Msh2-Msh3 MutS-homologs *(Surtees & Alani, 2006)* and Mus81 crossover-specific endonuclease *(Doe et al., 2004; Osman et al., 2003)* have been implicated in the Rad52-dependent GCR pathway *(Onaka et al., 2020; Onaka et al., 2016)*. As previously observed, *msh3*Δ and *mus81*Δ reduced GCRs in *rad51*Δ cells (Figure 4B). However, in *pcn1-K107R rad51*Δ cells, neither *msh3*Δ nor *mus81*Δ significantly reduced GCRs. These results demonstrate that Rad8-dependent PCNA K107 ubiquitination acts in the Rad52-dependent GCR pathway that involves Msh3 and Mus81 endonuclease.

**Figure 4.**
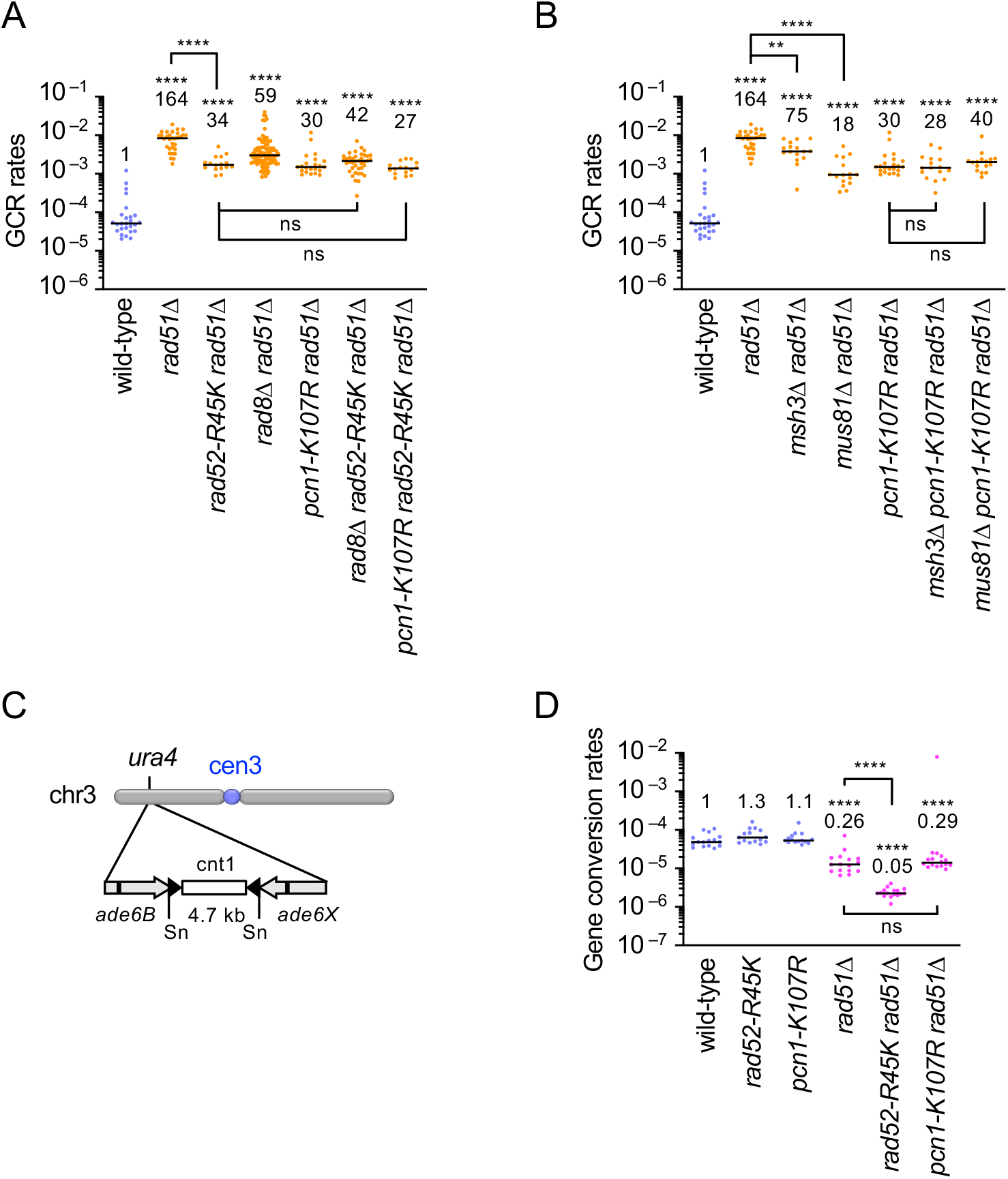
PCNA K107 is involved in the Rad52-dependent GCR pathway. (**A**) GCR rates of the wild-type, *rad51*Δ, *rad52-R45K rad51*Δ, *rad8*Δ *rad51*Δ, *pcn1-K107R rad51*Δ, *rad8*Δ *rad52-R45K rad51*Δ, and *pcn1-K107R rad52-R45K rad51*Δ strains (TNF5369, 5411, 6616, 5644, 6761, 6704, and 7006, respectively). (**B**) GCR rates of the wild-type, *rad51*Δ, *msh3*Δ *rad51*Δ, *mus81*Δ *rad51*Δ, *pcn1-K107R rad51*Δ, *msh3*Δ *pcn1-K107R rad51*Δ, and *mus81*Δ *pcn1-K107R rad51*Δ strains (TNF5369, 5411, 7081, 5974, 6761, 6990, and 7203, respectively). (**C**) A schematic diagram illustrates the *ade6B* and *ade6X* inverted repeats integrated at the *ura4* locus on the arm region of chr3. Sn, SnaBI. (**D**) Rates of gene conversion between *ade6B* and *ade6X* heteroalleles in the wild-type, *rad52-R45K, pcn1-K107R, rad51*Δ, *rad52-R45K rad51*Δ, and *pcn1-K107R rad51*Δ strains (TNF3631, 5995, 7837, 3635, 6021, and 7918 respectively). The two-tailed Mann-Whitney test. Non-significant (ns) *P* > 0.05; ** *P* < 0.01; **** *P* < 0.0001. Source data of the graphs are available in Figure 4—Source Data 1.

Rad52 promotes gene conversion as well as GCRs in the absence of Rad51 *(Onaka et al., 2020)*. However, Msh2-Msh3 and Mus81 are not required for the gene conversion branch of Rad52-dependent recombination. To determine whether PCNA K107 ubiquitination is also involved in gene conversion, we determined the rate of gene conversion between *ade6B* and *ade6X* heteroalleles that results in adenine prototrophs *(Zafar et al., 2017)* (Figure 4C, D). *rad52-R45K* reduced gene conversion in *rad51*Δ cells, as previously observed *(Onaka et al., 2020)*. However, *pcn1-K107R* did not reduce gene conversion. Together, these results indicate that, like Msh2-Msh3 and Mus81, PCNA K107 ubiquitination is specifically required for the GCR but not gene conversion branch of Rad52-dependent recombination.

### PCNA K107 ubiquitination weakens the interaction between PCNA subunits to facilitate GCRs

K107 is present in *C. elegans*, budding yeast, and fission yeast but not in humans, *M. musculus*, or *G. gallus* PCNA (Figure 5A). However, instead of K107, K110 is present in humans, *M. musculus*, and *G. gallus* but not in yeast PCNA. Structural analysis of fission yeast and budding yeast PCNA located K107 but not K164 at the interface between PCNA subunits (Figure 5B), suggesting that the ubiquitination of K107 weakens the interaction between PCNA subunits and changes the structure of the PCNA clamp. Interestingly, human K110 is also located at the interface, and its ubiquitination has been detected in cultured cells *(Kim et al., 2011)*, suggesting that human K110 is the counterpart of yeast K107. We reasoned that, if K107 ubiquitination weakens the interaction between PCNA subunits to cause GCRs, a mutation that impairs the interaction will bypass the requirement of K107 ubiquitination for GCRs. D150 is present at the PCNA-PCNA interphase (Figure 5B), and the D150E mutation has been shown to destabilize PCNA homotrimers *(Goellner et al., 2014; Johnson et al., 2016)*. Introduction of D150E into wild-type or *rad51*Δ cells resulted in small increases in GCR rates (Figure 5C). However, D150E dramatically increased GCR rates in *pcn1-K107R rad51*Δ cells to the level of *pcn1-D150E rad51*Δ. Most of the GCR products formed in *pcn1-K107R,D150E rad51*Δ cells were isochromosomes (Figure 5—figure supplement 1), showing that D150E bypassed the requirement of PCNA K107 for homology-mediated GCRs. D150E also bypassed Rad8 ubiquitin ligase requirement for GCRs (Figure 5D and Figure 5—figure supplement 1). D150E greatly increased GCR rates in *rad8-RING rad51*Δ cells. Together, these data show that Rad8-dependent PCNA K107 ubiquitination weakens the interaction between PCNA subunits to cause GCRs. Elg1 unloads PCNA from chromatin and facilitates recombination around stalled replication forks *(Kubota et al., 2015; Kubota et al., 2013; Tamang et al., 2019)*. If K107R mutation accumulates PCNA on DNA, which interferes with Rad52-dependent GCRs, loss of Elg1 will also reduce GCR rates. However, unlike PCNA K107R, *elg1*Δ did not significantly reduce GCRs in wild-type and *rad51*Δ cells (Figure 5E), indicating that Elg1-dependent PCNA unloading is not essential for GCRs.

**Figure 5.**
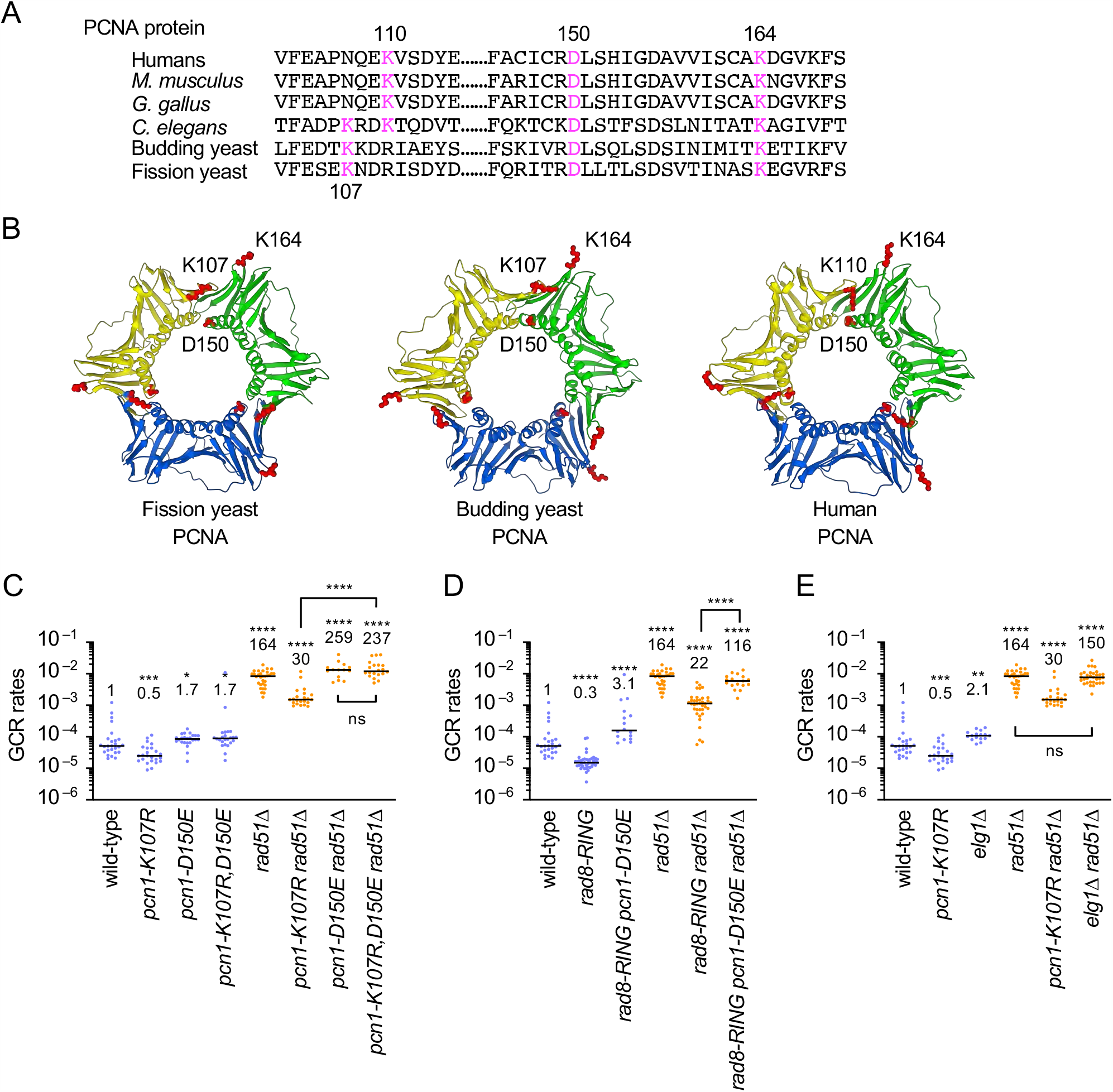
An interface mutation D150E bypasses the requirement of PCNA K107 ubiquitination for GCRs. (**A**) PCNA amino acid sequences that contain K107, K110, K150, or K164. (**B**) Structure of fission yeast PCNA was computed by the SWISS-MODEL program using an automated mode (Bienert et al., 2017). X-ray structure of fission yeast PCNA in a complex with an Spd1 peptide (PDB code 6qh1) was used as the template. X-ray structure of budding yeast PCNA (PDB code 1plr) and human PCNA (PDB code 1vym) are shown. Positions of PCNA K107, K110, D150, and K164 residues (red) were drawn using Mol* Viewer (https://molstar.org/). (**C**) GCR rates of the wild-type, *pcn1-K107R, pcn1-D150E, pcn1-K107R,D150E, rad51*Δ, *pcn1-K107R rad51*Δ, *pcn1-D150E rad51*Δ, and *pcn1-K107R,D150E rad51*Δ strains (TNF5369, 6738, 7724, 7727, 5411, 6761, 7744 and 7747, respectively). (**D**) GCR rates of the wild-type, *rad8-RING, rad8-RING pcn1-D150E, rad51*Δ, *rad8-RING rad51*Δ, and *rad8-RING pcn1-D150E rad51*Δ strains (TNF5369, 6207, 7750, 5411, 6219, and 7773, respectively). GCR products of *pcn1-K107R,D150E rad51*Δ *and rad8-RING pcn1-D150E rad51*Δ strains are shown in Figure 5—figure supplement 1. (**E**) GCR rates of the wild-type, *pcn1-K107R, elg1*Δ, *rad51*Δ, *pcn1-K107R rad51*Δ, and *elg1*Δ *rad51*Δ strains (TNF5369, 6738, 7696, 5411, 6761, and 7741, respectively). The two-tailed Mann-Whitney test. Non-significant (ns) *P* > 0.05; * *P* < 0.05; ** *P* < 0.01; *** *P* < 0.001; **** *P* < 0.0001. Source data of the graphs are available in Figure 5—Source Data 1.

## Discussion

Here, we found that fission yeast Rad8 facilitates Rad52-dependent GCRs through PCNA K107 ubiquitination. Loss of Rad8 reduced isochromosome formation in *rad51*Δ cells. Mutations in Rad8 HIRAN and RING finger but not helicase domain reduced GCRs. *mms2* and *ubc4* but not *ubc13* mutations; PCNA K107R but not K164R reduced GCRs in *rad51*Δ cells. The epistatic analysis showed that PCNA K107 ubiquitination plays a crucial role in the Rad52-dependent GCR pathway that involves Msh2-Msh3 and Mus81. PCNA K107 is located at the interface between PCNA subunits, suggesting that its ubiquitination interferes with the PCNA-PCNA interaction to cause GCRs. Indeed, an interface mutation D150E bypassed the requirement of PCNA K107 ubiquitination for GCRs. These data suggest that Rad8-dependent K107 ubiquitination changes the structure of the PCNA clamp to facilitate Rad52-dependent GCRs.

Our data suggest that Rad8 facilitates GCRs through 3’ DNA-end binding and ubiquitin ligase activity (Figure 6). Rad8 ubiquitin ligase acts with Mms2-Ubc13 ubiquitin-conjugating E2 enzymes to cause template switching *(Frampton et al., 2006)*. However, in the case of GCRs, Rad8 functions with Mms2-Ubc4, as *mms2* and *ubc4* but not *ubc13* mutations reduced GCRs in *rad51*Δ cells. That is further supported by the fact that *rad8-RING* and *ubc4* mutations epistatically reduce GCRs. Ubc4 and Ubc13 contain the cysteine residue critical for the E2 activity, while Mms2 is an E2 variant that lacks the active site *(VanDemark et al., 2001)*. Ubc13 but not Mms2 interacts with budding yeast Rad5 *(Ulrich & Jentsch, 2000)*. When Mms2-Ubc4 interacts with Rad8, Ubc4 rather than Mms2 may be the protein that directly binds Rad8 RING finger. E2 complexes determine substrate specificity for Rad8 ubiquitination: Mms2-Ubc4 promotes PCNA K107 while Mms2-Ubc13 causes PCNA K164 ubiquitination *(Das-Bradoo et al., 2010; Hoege et al., 2002)*. It is unknown how Rad8 choose the E2 partner. The helicase but not HIRAN domain is essential for PCNA K164 ubiquitination *(Achar et al., 2015; Ball et al., 2014; Choi et al., 2015)*. In contrast, the HIRAN but not helicase domain was required for GCRs. DNA-binding via the HIRAN domain may facilitate the RING finger to accommodate Mms2-Ubc4 rather than Mms2-Ubc13.

**Figure 6.**
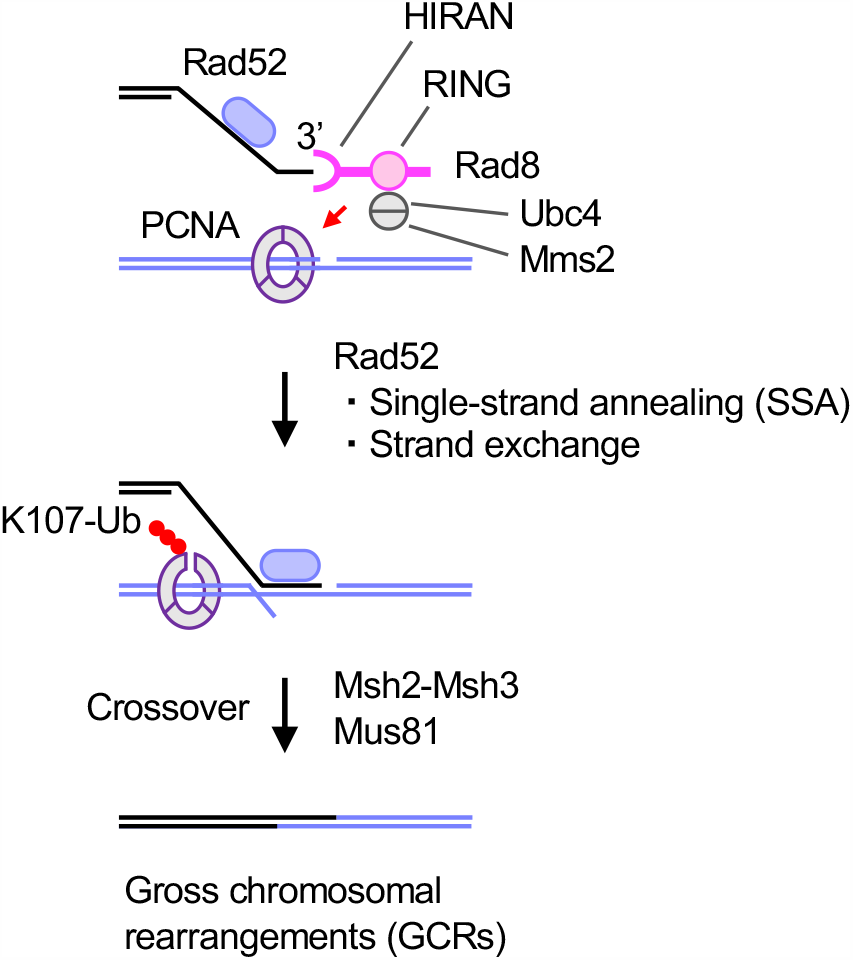
A model explains how Rad8 facilitates GCRs though PCNA K107 ubiquitination. Rad8 binds 3’ DNA-ends through HIRAN and interacts with Mms2-Ubc4 E2 complex through RING finger. The Rad8 complex ubiquitinates K107 of PCNA present on template DNA and weakens the interaction between PCNA subunits. The structural change of the PCNA clamp facilitates Rad52-dependent recombination and stimulates Msh2-Msh3 MutS-homologs and Mus81 endonuclease. Ub, ubiquitin.

Several lines of evidence suggest that Rad8-dependent PCNA K107 ubiquitination plays a crucial role in Rad52-dependent GCRs. Firstly, both Rad8 and Rad52 are involved specifically in homology-mediated GCRs. Like Rad52 *(Onaka et al., 2020)*, Rad8 was required for the formation of isochromosomes produced by recombination between centromere inverted repeats, but it was dispensable for chromosomal truncations that are produced by *de novo* telomere addition. Secondly, both *rad8*Δ and PCNA K107R reduced GCRs epistatically with the *rad52-R45K* mutation that impairs SSA activity *(Onaka et al., 2020)*. Thirdly, PCNA K107R reduced GCRs epistatically with loss of Msh3 or Mus81, both of which have been implicated in Rad52-dependent GCRs *(Onaka et al., 2020)*. Rad52 facilitates gene conversion between inverted repeats in the absence of Rad51. However, like Msh2-Msh3 and Mus81, PCNA K107 was dispensable for the gene conversion in *rad51*Δ cells, indicating that PCNA K107 ubiquitination is specifically involved in the GCR branch of Rad52-dependent recombination.

Post-translational modifications of PCNA at K164 affect protein interaction. K164 ubiquitination facilitates PCNA binding to translesion synthesis polymerases and Mgs1/ZRANB3 helicase *(Leung et al., 2018)*. In budding yeast, Small ubiquitin-like modifier (SUMO) protein is also attached to PCNA K164. K164 SUMOylation causes PCNA interaction with Srs2 helicase that suppresses Rad51-dependent recombination by dissociating Rad51 protein from ssDNA *(Pfander et al., 2005)*. However, the C-terminal domain of budding yeast Srs2 that interacts with SUMOylated PCNA is not conserved in fission yeast Srs2 *(Frampton et al., 2006)*. Nonetheless, loss of Rad51 resulted in slow growth phenotypes and increased sensitivities in DNA damaging agents in *pcn1-K164R* but not in *pcn1-K107R* cells *(Frampton et al., 2006)* (Figure 3—figure supplement 3). PCNA K164R slightly but significantly reduced GCRs in wild-type but not in *rad51*Δ cells (Figure 3C). PCNA K164R may channel DNA repair into Rad51-dependent gene conversion pathway.

How does PCNA K107 ubiquitination cause GCRs? Our data suggest that K107 ubiquitination changes the PCNA clamp structure to facilitate GCRs (Figure 6). PCNA subunits interact with each other to form ring-shaped homotrimers. K107 is located at the interface between PCNA subunits. An interface mutation D150E *(Goellner et al., 2014; Johnson et al., 2016)* bypassed the requirement of Rad8 RING finger and PCNA K107 for GCRs, showing that the ubiquitin or ubiquitin chain at K107 is not essential for GCRs and that weakening the PCNA-PCNA interaction is sufficient to cause GCRs. PCNA K107R mutation did not significantly change the level of Rad52 foci accumulated in the absence of Rad51 *(Miyazaki et al., 2004)* (Figure 3—figure supplement 4), suggesting that PCNA K107 ubiquitination is not required for Rad52 to localize to damage sites. However, the PCNA clamp present at the end of Okazaki fragments can be a structural hindrance for Rad52-dependent SSA or strand exchange reaction. The structural change induced by K107 ubiquitination might facilitate translocation or dissociation of the PCNA clamp away from Okazaki fragment ends, thereby promoting assembly of recombination enzymes on small ssDNA gaps between Okazaki fragments (Figure 6). Consistent with this idea, in budding yeast DNA ligase I mutants, K107 ubiquitination facilitates checkpoint activation that depends on the assembly of checkpoint proteins onto ssDNA *(Das-Bradoo et al., 2010; Nguyen et al., 2013)*. Alternatively, the structural change induced by K107 ubiquitination might recruit Msh2-Msh3 or Mus81 to the PCNA clamp to promote GCR reaction (Figure 6). Like Msh2-Msh3 MutS-homologs and Mus81 endonuclease *(Onaka et al., 2020)*, PCNA K107 ubiquitination is specifically required for the GCR branch of Rad52-dependent recombination. It has been reported that the PCNA clamp can interact with Msh2-Msh3 *(Clark et al., 2000)* and Mus81 to enhance endonuclease activity *(Sisakova et al., 2017)*.

RFC-like complexes containing Elg1 unload PCNA from DNA after Okazaki fragment ligation *(Kubota et al., 2015; Kubota et al., 2013)*. Elg1 promotes Rad51 and Rad52 proteins’ recruitment to stalled replication forks and facilitates nearby recombination *(Tamang et al., 2019)*, suggesting that PCNA unloading facilitates recombination around stalled forks. An interface mutation D150E also bypasses the role of Elg1 in PCNA unloading and recombination around stalled replication forks *(Johnson et al., 2016; Tamang et al., 2019)*. However, unlike PNCA K107R, loss of Elg1 did not reduce GCRs, indicating that Elg1-dependent PNCA unloading is not essential for GCRs. We do not exclude the possibility that K107 ubiquitination causes PCNA unloading, by itself or with the factors other than Elg1.

This study has uncovered the role of Rad8-dependent PCNA K107 ubiquitination in Rad52-dependent GCRs. We also suggest that human PCNA K110 is the counterpart of yeast PCNA K107. Interestingly, HLTF, the human homolog of Rad8, is often amplified and overexpressed in cancer *(Bryant et al., 2019)*, suggesting that HLTF causes GCRs and facilitates tumorigenesis.

## Materials and Methods

### Genetic procedures and yeast strains

The fission yeast strains used in this study are listed in Supplementary File 1. Standard genetic procedures and fission yeast media were used as previously described *(Onaka et al., 2016)*. Pombe minimal glutamate (PMG) medium is identical to Edinburgh minimal medium (EMM), except containing 3.75 g/l monosodium L-glutamate instead of 5 g/l ammonium chloride. Yeast nitrogen base (YNB) medium contained 1.7 g/l yeast nitrogen base (BD Biosciences, San Jose, California, Difco 233520), 5 g/l ammonium sulphate (Nacalai Tesque, Kyoto, Japan, 02619-15), and 2% glucose. YE, YNB, EMM, and PMG contain 225 mg/l of each amino acid when indicated. FOA+UA is a YNB derivative supplemented with 56 mg/l uracil, 225 mg/l adenine, and 1 g/l 5-fluoroorotic acid monohydrate (Apollo Scientific, Stockport, United Kingdom, PC4054). Yeast cells were grown at 30°C.

The *rad8-HIRAN* mutant strain was created by the pop-in/pop-out gene replacement *(Grimm et al., 1988)*. pTN1192 plasmid containing *ura4*^+^ and *rad8-HIRAN* was linearised by BglII digestion at a unique site in the *rad8* region and introduced into *ura4-D18* cells. Ura^+^ transformants were selected on EMM. After confirmation of the correct integration by PCR and DNA sequencing, the *rad8*:*ura4*^+^:*rad8-HIRAN* strain was grown in YE media supplemented with uracil (YE+U) and then plated on FOA+U media to select Ura^−^ colonies, resulting from “pop-out” of the *ura4*^+^ marker. DNA sequencing confirmed the retention of the *rad8*-*HIRAN* mutation in the Ura^−^ cells. *rad8-Helicase* and *rad8-RING* mutant strains were created essentially in the same way, but BglII-digested pTN1191 plasmid containing *rad8-Helicase* and PacI-digested pTN1193 plasmid containing *rad8-RING* were used, respectively.

The *pcn1-K107R* mutant strain was constructed as follows. First, we created the DNA fragment containing the *pcn1-K107R* mutation. Two partially overlapping fragments were amplified separately from yeast genomic DNA: a 0.6 kb fragment using pcn1-F3 and pcn1-K107RB primers, and a 1.4 kb fragment using pcn1-K107RT and pcn1-R2 primers. pcn1-K107RB and pcn1-K107RT contain the *pcn1-K107R* mutation. The two PCR products were mixed and connected by the second round of PCR in the presence of pcn1-F3 and pcn1-R2 primers, resulting in the formation of a 1.9 kb fragment containing the *pcn1-K107R* mutation. To create the *pcn1-K107R* strain, we first introduced the *ura4*^+^ gene 0.7 kb downstream of the *pcn1*^+^ gene in *ura4-D18* cells, making *pcn1*^+^:*ura4*^+^ cells. Then, the 1.9 kb PCR fragment that contains the *pcn1-K107R* mutation and encompasses the *ura4*^+^ integration site was introduced into *pcn1*^+^:*ura4*^+^ cells. Ura^−^ transformants were selected on FOA+U plates. DNA sequencing confirmed the correct integration of *pcn1-K107R. pcn1-K107A, pcn1-K164R*, and *pcn1-D150E* strains were created essentially in the same way, except that pcn1-K107AB/pcn1-K107AT, pcn1-K164RB/pcn1-K164RT, and pcn1-D150EB/pcn1-D150ET primers, instead of pcn1-K107RB/pcn1-K107RT, were used to create *pcn1-K107A, pcn1-K164R*, and *pcn1-D150E* mutations, respectively. The *pcn1-K107R,D150E* double mutation was created using genomic DNA of *pcn1-K107R* cells and pcn1-D150EB/pcn1-D150ET primers.

We created the *ubc4-P61S* mutation as follows. Two partially overlapping fragments were amplified separately: a 0.7 kb fragment using ubc4-F1 and ubc4-P61S-B primers, and a 1.5 kb fragment using ubc4-P61S-T and ubc4-R4 primers. ubc4-P61S-B and ubc4-P61S-T contain the *ubc4-P61S* mutation. The two PCR products were mixed and connected by the second round of PCR in the presence of ubc4-F1 and ubc4-R4 primers, resulting in the formation of a 2.2 kb fragment containing the *ubc4-P61S* mutation. To create the *ubc4-P61S* strain, we first introduced the *ura4*^+^ gene 0.6 kb downstream of the *ubc4*^+^ gene in *ura4-D18* cells, making *ubc4*^+^:*ura4*^+^ cells. Then, the 2.2 kb PCR fragment that contains the *ubc4-P61S* mutation and encompasses the *ura4*^+^ integration site was introduced into *ubc4*^+^:*ura4*^+^ cells. Ura^−^ transformants were selected on FOA+U plates. DNA sequencing confirmed the correct integration of *ubc4-P61S*. Sequences of PCR primers used are listed in Supplementary File 2.

### Plasmids

We constructed the plasmid pTN1192 containing *ura4*^+^ and *rad8-HIRAN* as follows. Two partially overlapping fragments were amplified separately from yeast genomic DNA: a 1.4 kb fragment using rad8-KpnI and rad8-HIRAN-B primers, and a 1.0 kb fragment using rad8-HIRAN-T and rad8-SalI primers. rad8-HIRAN-B and rad8-HIRAN-T contain the *rad8-HIRAN* mutation. The two PCR products were mixed and connected by the second round of PCR in the presence of rad8-KpnI and Rad8-SalI primers. A 2.3 kb KpnI-SalI restriction fragment of the PCR product containing the *rad8-HIRAN* mutation was introduced between KpnI-SalI sites of pTN782 *(Okita et al., 2019)*, which contains a 1.5 kb HindIII-SspI fragment containing the *ura4*^+^ gene between HindIII-EcoRV sites of pBluescript II KS^+^ (Agilent, Santa Clara, California).

The plasmid pTN1191 containing *ura4*^+^ and *rad8-Helicase*, and the plasmid pTN1193 containing *ura4*^+^ and *rad8-RING*, were created essentially in the same way as described above. To create pTN1191, rad8-SacII/rad8-Helicase-B/rad8-Helicase-T/rad8-BamHI primers were used, and a 2.9 kb SacII-BamHI restriction fragment of the 2nd PCR product was introduced between SacII-BamHI sites of pTN782. To create pTN1193, rad8-1/rad8-RING-B/rad8-RING-T/rad8-EcoRI primers were used, and a 1.1 kb BamHI-EcoRI restriction fragment of the 2nd PCR product was introduced between BamHI-EcoRI sites of pTN782.

### GCR rates

Spontaneous rates of GCRs that result in loss of *ura4*^+^ and *ade6*^+^ markers were determined essentially as previously described *(Onaka et al., 2020)*. Yeast cells harbouring ChL^C^ were incubated on EMM+UA plates for 6–8 days. 10 ml EMM+UA was inoculated with a single colony from EMM+UA plates and incubated for 1–2 days. Cells were plated on YNB+UA and FOA+UA plates and incubated for 5–8 days. Leu^+^ and Leu^+^ Ura^−^ colonies formed on YNB+UA and FOA+UA plates, respectively, were counted. Leu^+^ Ura^−^ colonies were streaked on EMM+UA plates to examine the colony size and transferred to EMM+U plates to inspect adenine auxotrophy. The number of Leu^+^ Ura^−^ Ade^−^ was obtained by subtracting the number of Leu^+^ Ura^−^ Ade^+^ from that of Leu^+^ Ura^−^. Rates of GCR per cell division were calculated *(Lin et al., 1996)*, using the numbers of Leu^+^ cells and Leu^+^ Ura^−^ Ade^−^ cells, using Microsoft Excel for Mac 16 (Microsoft, Redmond, Washington).

### Pulsed-field gel electrophoresis (PFGE) analysis of GCR products

1×10^8^ cells grown at 25°C were collected, suspended in 2.5 ml ice-cold 50 mM EDTA, and stored at 4°C. After centrifugation, cells were suspended in 1 ml CSE buffer (20mM citrate phosphate, 1 M sorbitol, 50 mM EDTA, pH 5.6). After adding 5 to 10 µl Zymolyase 20T (Seikagaku, Tokyo, Japan, 25 mg/ml) and 5 to 10 µl lyzing enzyme (Sigma, St. Louis, Missouri, 25 mg/ml), cell suspensions were incubated for 20 to 50 min at 30°C. After centrifugation at 700 rpm for 10 min at 4°C (TOMY, Tokyo, Japan, MX-201, TMS-21 swing rotor), the pellet was suspended in 140 µl CSE buffer. After adding 140 µl 1.6% low melting agarose gel (Nacalai Tesque, 01161-12) pre-heated at 50°C, the suspension of spheroplasts was transferred into a mould. After 20 min at 4°C, the agarose plugs were incubated in SDS-EDTA solution (1% SDS, 0.25 M EDTA) for 2 h at 60°C. The plugs were transferred into ESP solution (0.5M EDTA, 1% N-lauroylsarcosine, 1.5 mM calcium acetate) supplemented with 0.5 mg/ml proteinase K and incubated overnight at 50°C. The plugs were transferred into another ESP solution supplemented with 0.5 mg/ml proteinase K and incubated at 50°C for an additional 8 h. The plugs were stored in TE buffer (10 mM Tris-HCl (pH8.0), 1 mM EDTA) at 4°C. Chromosomal DNAs prepared in agarose plugs were resolved using a CHEF-DRII pulsed-field electrophoresis system (Bio-Rad, Hercules, California). Broad-range PFGE ran at 2 V/cm with a pulse time of 1500 s or 1600 s for 42 h followed by 2.4 V/cm with a pulse time of 180 s for 4 h, at 4°C in 1×TAE buffer (40 mM Tris-acetate, 1 mM EDTA) using 0.55% Certified Megabase agarose gel (Bio-Rad, 161-3109). Short-range PFGE ran at 4.2 or 4.5 V/cm with a pulse time from 40 to 70 s for 24 h, at 4°C in 0.5×TBE buffer (89 mM Tris-borate, 2 mM EDTA) using 0.55% Certified Megabase agarose gel, otherwise indicated. DNA was stained with 0.2 µg/ml ethidium bromide (EtBr) (Nacalai Tesque, 14631-94) and detected using a Typhoon FLA9000 gel imaging scanner (GE Healthcare, Chicago, Illinois). Gel images were processed using ImageJ 1.8.0 (NIH, United States).

### Gene conversion rates

Spontaneous rates of gene conversion that occurs between *ade6B* and *ade6X* heteroalleles integrated at the *ura4* locus *(Zafar et al., 2017)* were determined. 10 ml PMG+UA was inoculated with a single colony from PMG+UA plates and incubated for 1–2 days. Cells were plated onto PMG+UA and PMG+U media. After 4–7 days’ incubation of the plates, colonies formed on PMG+UA and PMG+U were counted to determine the number of colony-forming units and Ade^+^ prototrophs, respectively. The rates of gene conversion per cell division were calculated *(Lin et al., 1996)*, using Microsoft Excel for Mac 16.

### Cell imaging

Exponentially growing cells in EMM were collected, seeded on glass-bottom dishes (Matsunami Glass, Osaka, Japan, D11130H), and observed using a DeltaVision Personal fluorescence microscopy system (GE Healthcare), which is based on an Olympus wide-field IX71 fluorescence microscope equipped with a CoolSNAP HQ2 CCD camera (Photometrics, Tucson, Arizona) and an oil-immersion objective lens (UAPO 40×; NA = 1.35; Olympus, Tokyo, Japan).

### Statistical analyses

Two-tailed Mann-Whitney tests and Fisher’s exact tests were performed using GraphPad Prism for Mac version 8 (GraphPad Software, San Diego, California). Two-tailed student’s *t*-tests were performed using Microsoft Excel for Mac 16.

## Acknowledgements

We thank Crystal Tang, Keitaro Oki, and Hirofumi Ohmori (Osaka University) for technical assistance, Yumiko Kubota, Akiko Okita, Atsushi Onaka, Crystal Tang, Xu Ran, and Piyusha Mongia (Osaka University) for comments on the manuscript. We also thank Anthony M. Carr (University of Sussex), Jürg Bähler (University College London), and Aki Hayashi (National Institutes for Basic Biology) for providing plasmids.

---

**Figure 1—figure supplement 1.**
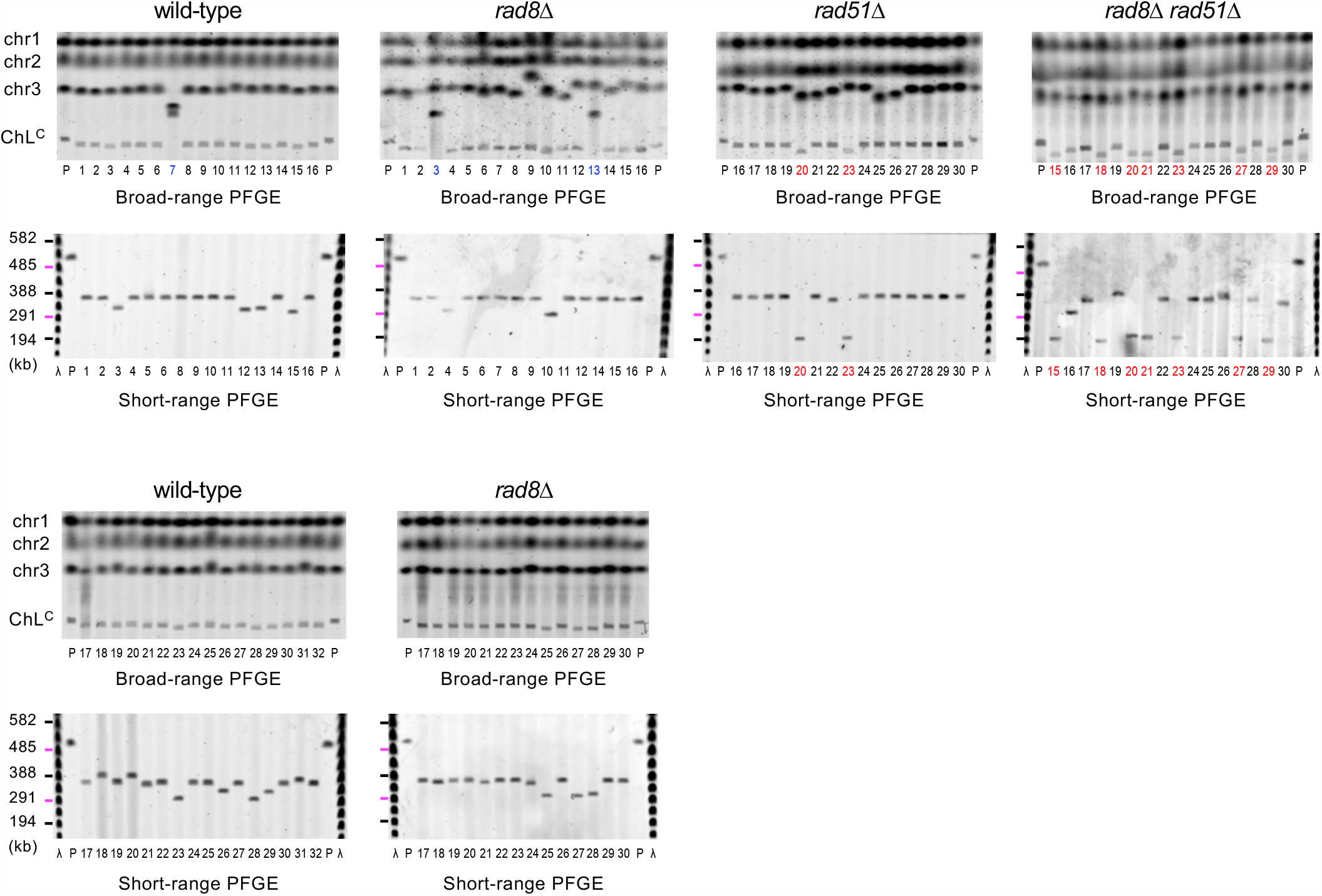
GCR products formed in wild-type, *rad8*Δ, *rad51*Δ and *rad8*Δ *rad51*Δ cells. Chromosomal DNAs prepared from the parental and independent GCR clones of the wild-type, *rad8*Δ, *rad51*Δ and *rad8*Δ *rad51*Δ strains (TNF5369, 5549, 5411, and 5644, respectively) were separated by broad- and short-range PFGE and stained with EtBr. Sample number of translocations and truncations are highlighted in blue and red, respectively. Uncropped images of the gels are available in Figure 1— Source Data 2.

**Figure 3—figure supplement 1.**
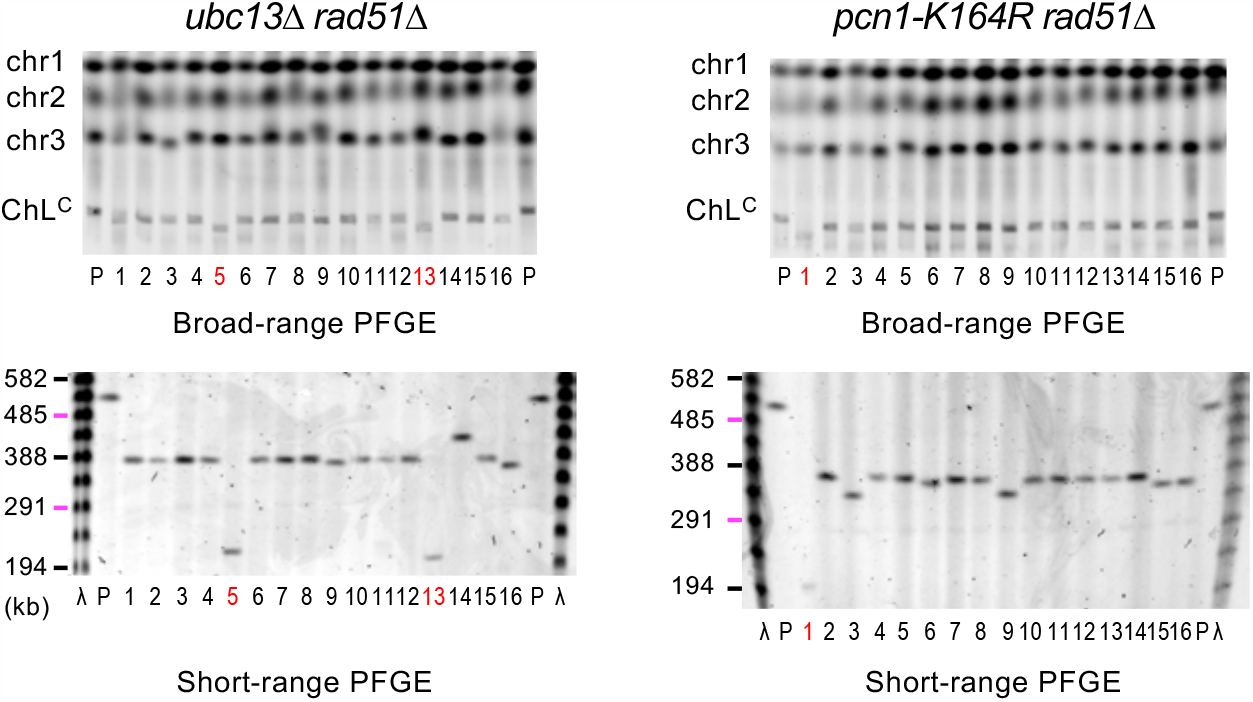
GCR products formed in *ubc13*Δ *rad51*Δ and *pcn1-K164R rad51*Δ cells. Chromosomal DNAs prepared from the parental and independent GCR clones of the *ubc13*Δ *rad51*Δ and *pcn1-K164R rad51*Δ strains (TNF6115 and 6104, respectively) were separated by broad- and short-range PFGE and stained with EtBr. Short-range PFGE ran at 4.5 V/cm with a pulse time from 4 to 120 s for 48 h at 4°C. Uncropped images of the gels are available in Figure 3—Source Data 2.

**Figure 3—figure supplement 2.**
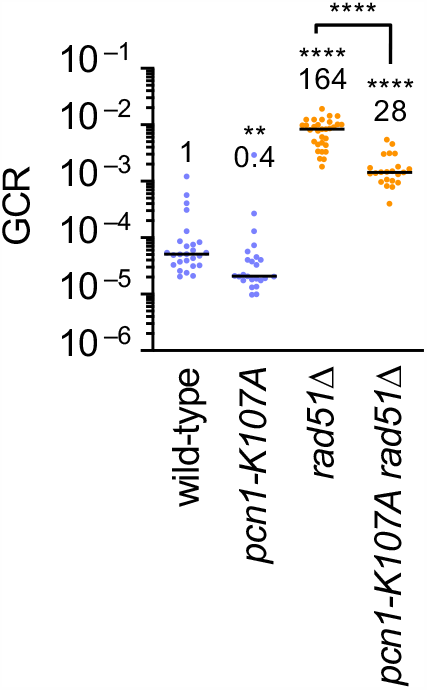
The *pcn1-K107A* mutation reduces GCR rates. GCR rates of the wild-type, *pcn1-K107A, rad51*Δ, *pcn1-K107A rad51*Δ strains (TNF5369, 6699, 5411, and 6719, respectively). The two-tailed Mann-Whitney test. ** *P* < 0.01; **** *P* < 0.0001. Source data are available in Figure 3— Source Data 1.

**Figure 3—figure supplement 3.**
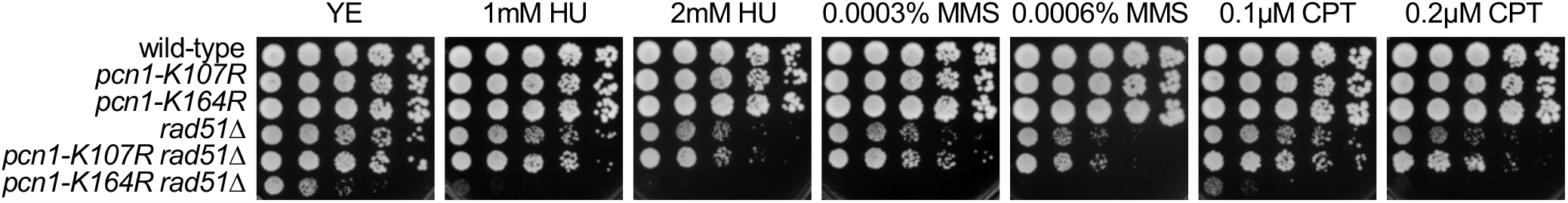
DNA damage sensitivity of the wild-type, *pcn1-K107R, pcn1-K164R, rad51*Δ, *pcn1-K107R rad51*Δ, and *pcn1-K164R rad51*Δ strains (TNF35, 6968, 6948, 2610, 6988, and 6986, respectively). Exponentially growing cells in YE media were 5-fold serially diluted and spotted onto YE plates supplemented with hydroxyurea (HU), methyl methanesulfonate (MMS), or camptothecin (CPT) at the indicated concentrations.

**Figure 3—figure supplement 4.**
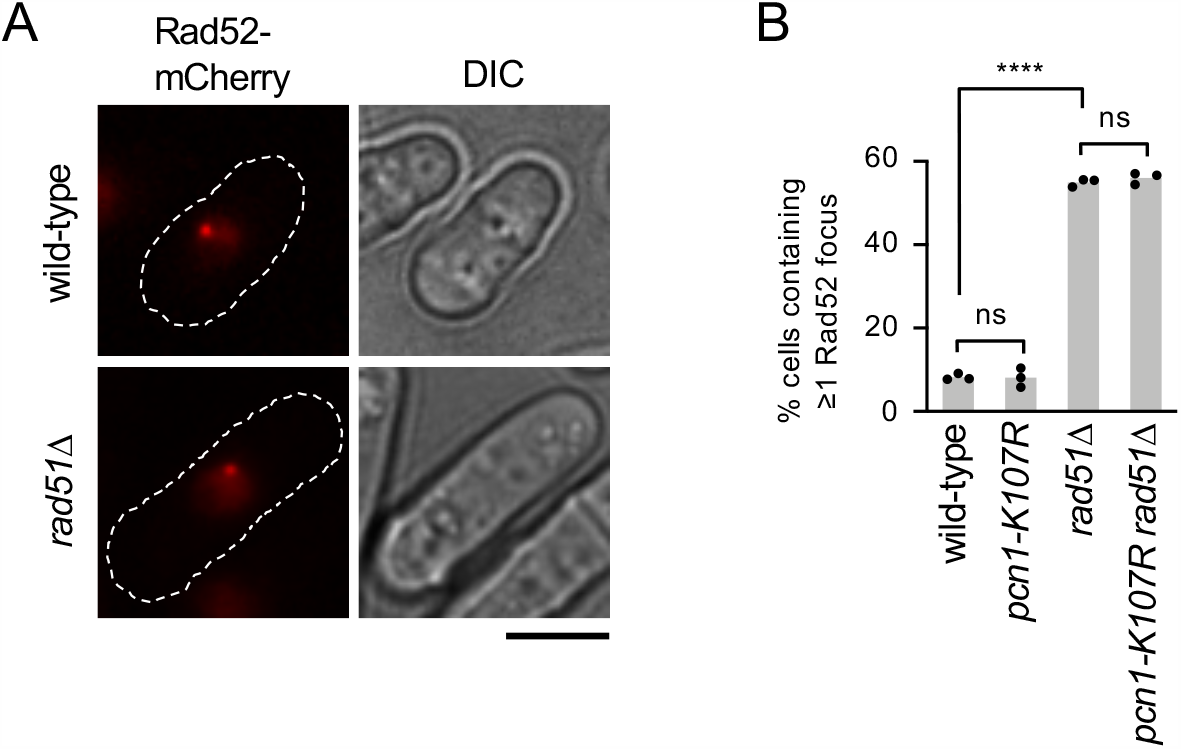
PCNA K107 is not essential for Rad52 focus formation. (**A**) Rad52-mCherry foci were observed by fluorescence microscopy. DIC, differential interference contrast. A scale bar indicates 5 μm. (**B**) Percentages of cells containing Rad52 foci in the wild-type, *pcn1-K107R, rad51*Δ, and *pcn1-K107R rad51*Δ strains (TNF4462, 7387, 7800, and 7802, respectively). Bars represent the mean of three independent experiments shown as dots. > 200 cells were counted in each experiment. The two-tailed student’s *t*-test. Non-significant (ns) *P* > 0.05; **** *P* < 0.0001. Source data of the graph are available in Figure 3–Source Data 1.

**Figure 5—figure supplement 1.**
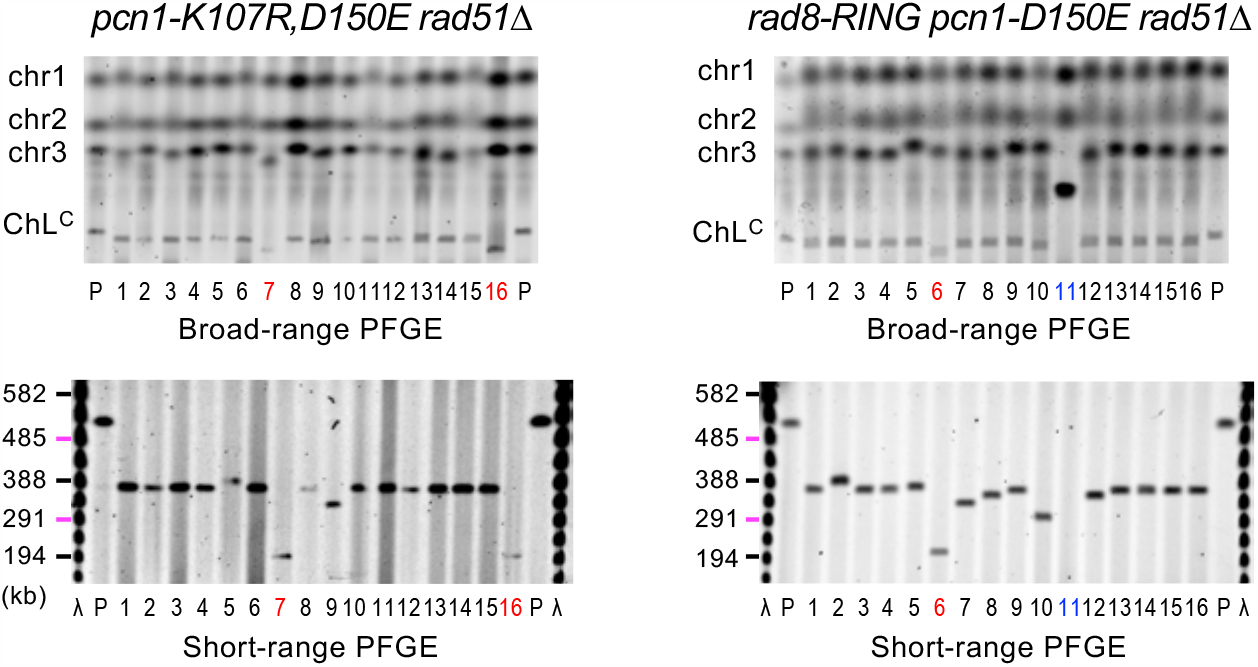
GCR products formed in *pcn1-K107R,D150E rad51*Δ and *rad8-RING pcn1-D150E rad51*Δ cells. Chromosomal DNAs prepared from the *pcn1-K107R,D150E rad51*Δ and *rad8-RING pcn1-D150E rad51*Δ strains (TNF7747 and 7773, respectively) were separated by broad- and short-range PFGE and stained with EtBr. Uncropped images of the gels are available in Figure 5—Source Data 2.

